# Chromosome painting does not support a sex chromosome turnover in Lacerta agilis Linnaeus, 1758

**DOI:** 10.1101/846501

**Authors:** Artem P. Lisachov, Massimo Giovannotti, Jorge C. Pereira, Daria A. Andreyushkova, Svetlana A. Romanenko, Malcolm A. Ferguson-Smith, Pavel M. Borodin, Vladimir A. Trifonov

## Abstract

Reptiles show a remarkable diversity of sex determination mechanisms and sex chromosome systems, derived from different autosomal pairs. The origin of the ZW sex chromosomes of *Lacerta agilis*, a widespread Eurasian lizard species, is a matter of discussion: is it a small macrochromosome from the 11-18 group, common to all lacertids, or this species has unique ZW pair derived from the large chromosome 5. Using independent molecular cytogenetic methods, we investigated the karyotype of *L. agilis exigua* from Siberia, Russia, to identify the sex chromosomes. FISH with the flow-sorted chromosome painting probe, derived from *L. strigata* and specific to chromosomes 13, 14, and Z, confirmed that the Z chromosome of *L. agilis* is a small macrochromosome, the same as in *L. strigata*. FISH with the telomeric probe showed an extensive accumulation of the telomeric repeat on the W chromosome in agreement with previous studies, excluding the possibility that the lineages of *L. agilis* studied in different works could have different sex chromosome systems due to a putative intra-species polymorphism. Our results reinforce the idea of the stability of the sex chromosomes and lack of evidence for sex-chromosome turnovers in known species of Lacertidae.

## Introduction

Reptiles have diverse mechanisms of sex determination. In some taxa, such as crocodiles, red-eared turtles, and leopard geckos, the sex of the offspring is determined by the temperature of egg incubation, and there are no specific sex-linked genes or chromosomes [Viets et al., 1993]. In other lineages, different chromosomes independently acquired sex-determining genes (via mutations of their original genes or translocations of genes from other chromosomes), and became sex chromosomes of either XX/XY or ZZ/ZW type [Pokorná and Kratochvíl, 2009]. The most studied reptilian sex chromosomes are those of pleurodont iguanas (XX/XY) [Alföldi et al., 2011; Gamble et al., 2014; Rovatsos et al., 2014; Kichigin et al., 2016; Giovannotti et al., 2017; Lisachov et al., 2019], advanced snakes (ZZ/ZW) [Matsubara et al., 2006; Vicoso et al., 2013; Rovatsos et al., 2015], anguimorphs (ZZ/ZW) [Matsubara et al., 2014; Rovatsos et al., 2019a]. In certain cases, sex chromosome systems are not universal for such large clades, but are specific to one species or a group of related species. This is a rather common situation in geckos [Gamble, 2010].

For all reptile sex chromosome systems, the exact master sex-determining genes and therefore the molecular mechanisms underlying sex determination are unknown. Uncovering these mechanisms will advance the understanding of general principles of the evolution of sex determination. It may also help to predict the sensitivity of sex chromosome systems to climate change. Extreme temperatures can override chromosomal sex determination in some squamate reptiles (described for Agamidae and Scincidae), skew the sex ratio and thus make their populations unstable and prone for extinction [Holleley et al., 2015].

The first step in finding the master sex determining genes is uncovering the correspondence between a certain reptile sex chromosome system and the homologous chromosome or syntenic block in the reference sauropsid genomes, such as chicken (*Gallus gallus*, GGA) and anole lizard (*Anolis carolinensis*, ACA). For example, the XY chromosomes of *Anolis* correspond to the chicken chromosome 15 (GGA15) [Alföldi et al., 2011]. This “genomic identity” is known for many reptile sex chromosome systems, and can be uncovered by different molecular genetic, genomic, and cytogenetic approaches [Deakin and Ezaz, 2019].

However, for some taxa different works and methods produced contradicting results. This can be due to intraspecific polymorphisms, wrong species identifications or technical errors (discussed by Gamble, [2010]). The sand lizard (*Lacerta agilis*), a common Eurasian species, is one of the examples of such contradiction. It belongs to the family Lacertidae, which is long-known to have a ZW sex chromosome system [Ivanov and Fedorova, 1970].

The lacertid chromosomes generally share the same acrocentric morphology and gradually decrease in length, therefore individual chromosomes are difficult to distinguish. Although the W chromosome can be identified by heterochromatinization and repetitive sequence accumulation [Capriglione et al., 1994], the Z chromosome is “hidden” among the autosomes, and its genetic content and even the position in the karyotype remained unknown for a long time.

Srikulnath et al. [2014] putatively identified the chromosome 5 of *L. agilis* from Sweden (*L. agilis agilis*) (LAG5) as the Z chromosome using Hoechst staining. Using FISH mapping of cDNA probes of protein-coding genes, they found that this chromosome is a homologue of the short arm of the *Anolis carolinensis* chromosome 3 (ACA3p) and of the chicken chromosomes 6 and 9 (GGA6, GGA9). In another work, Matsubara et al. [2015] described the W chromosome of these lizards, and found that it contains C-positive heterochromatin, enriched with telomere-like sequences.

Later, Rovatsos et al. [2016; 2019b] used transcriptome analysis and qPCR to identify hemizygous and thus suggestively Z-linked genes in several species of lacertids, including *L. agilis* from the Czech Republic (*L. agilis argus*). In these works, the genes identified by Srikulnath et al. [2014] as Z-specific were identified as (pseudo)autosomal. The homeologues of the Z-linked genes which were identified by Rovatsos et al. [2016, 2019b] are located in two microchromosomes of *A. carolinensis*: ACA11 and ACA16 (homologous to GGA4p and GGA17, respectively) [Kichigin et al., 2016]. The authors suggested that the Z chromosome of lacertids should be one of the small macrochromosomes, formed via fusion of two ancestral squamate microchromosomes [Uno et al., 2012]. Their results were supported by the genome sequencing project of *Podarcis muralis* [Andrade et al., 2019].

There are two possible explanations for this contradiction. First, it is possible that the different lineages of *L. agilis*, studied in these two works, indeed have different sex chromosome systems. Namely, the lineage studied by Srikulnath et al. [2014] might have experienced a sex chromosome turnover, leading to the loss of the original lacertid sex chromosome system and appearance of a new system, based on another chromosome pair. The cases of such turnovers, when a taxon “forsakes” a well-established sex chromosome system and acquires a new one, are rather rare [Pokorná and Kratochvíl, 2016], but known in lizards [Nielsen et al., 2019] and even in mammals [Matveevsky et al., 2017]. An intraspecific polymorphism in sex-determining systems is also possible, as in the case of the frog *Glandirana rugosa* [Ogata et al., 2018]. The second possibility is that one or both these identifications of the *L. agilis* sex chromosome are erroneous.

No additional studies with independent methods were conducted so far to investigate this issue. Meanwhile, resolving this contradiction is crucial for understanding the patterns of evolution of sex-determining mechanisms in reptiles in general. If confirmed, “the curious case of *Lacerta agili*s” could become an interesting model to study the processes of sex determination evolution.

In this work, we investigated the karyotype of *L. agilis exigua* from Novosibirsk (Russia) to identify and describe its Z and W chromosomes. The Z chromosome was identified using FISH with a flow-sorted chromosome painting probe derived from *L. strigata*, containing its chromosomes 13, 14 and Z. Sorted chromosome probes of lacertids were generated and chromosome painting was conducted previously, but with limited success, and *L. agilis* and *L. strigata* were not studied [Rojo Oróns, 2015]. The W-chromosome was identified by its size and using FISH with a telomere-specific probe.

## Materials and methods

The fibroblast cell culture of *L. strigata*, here used for flow-sorting, was established previously by Giovannotti et al. [2018]. The flow-sorted chromosome libraries were obtained using a Mo-Flo® (Beckman Coulter) high-speed cell sorter at the Cambridge Resource Centre for Comparative Genomics, Department of Veterinary Medicine, University of Cambridge, Cambridge, UK, as described previously [Yang et al., 1995]. The painting probes were generated from the DOP-PCR amplified libraries by a secondary DOP-PCR incorporation of Flu12-dUTP (Bioron, Germany) [Telenius et al., 1992].

The fibroblast cell culture of *L. agilis* was obtained from the muscle tissues of one juvenile female *L. agilis exigua*, originating from Novosibirsk province (Russia), in the Laboratory of Animal Cytogenetics, the Institute of Molecular and Cellular Biology, Russia, using enzymatic treatment of tissues as described previously [Stanyon and Galleni 1991, Romanenko et al. 2015]. All cell lines were deposited in the IMCB SB RAS cell bank (“The general collection of cell cultures”, 0310-2016-0002). Metaphase chromosome spreads were prepared from chromosome suspensions obtained from early passages of primary fibroblast cultures as described previously [Yang et al. 1999, Graphodatsky et al. 2000, 2001].

The telomere-specific probe was generated using PCR with self-annealing oligonucleotide primers, as described previously [Ijdo et al., 1991], and labelled with TAMRA-dUTP (Biosan, Novosibirsk, Russia) in a secondary PCR. FISH was performed with standard techniques [Liehr et al., 2017]. The preparations were analyzed with an Axioplan 2 Imaging microscope (Carl Zeiss) equipped with a CCD camera (CV M300, JAI), CHROMA filter sets, and the ISIS4 image processing package (MetaSystems GmbH). The brightness and contrast of all images were enhanced using Corel PaintShop Photo Pro X6 (Corel Corp).

## Results

The karyotype of *L. strigata* consisted of 38 chromosomes with a heteromorphic W chromosome, in agreement with previous descriptions (Fig. 1a) [Ivanov and Fedorova, 1970]. Here, for the first time, we have obtained flow-sorted chromosome libraries of a *Lacerta* species. The sorted karyotype of *L. strigata* was comprised of 17 peaks (LST_A to LST_Q) (Fig. 2). The probe derived from the peak LST_L was found to hybridize to five chromosomes on the metaphases of the same specimen used for establishing the fibroblast cell culture. Therefore, it was concluded that this peak contained two autosome pairs and the Z chromosome (Fig. 3a). The absence of the signal on the W chromosome is apparently due to the fact that the W chromosome of lacertids is too degenerated and does not hybridize with the DNA from the Z chromosome [Rovatsos et al., 2019b; Andrade et al., 2019]. The autosomes were identified as pairs 13 and 14 basing on the position of the LST_L peak in the flow karyotype. The co-occurrence of several chromosomes in the same probe is due to their very similar size and GC-content, which did not allow to separate them during chromosome sorting.

**Fig. 1.**
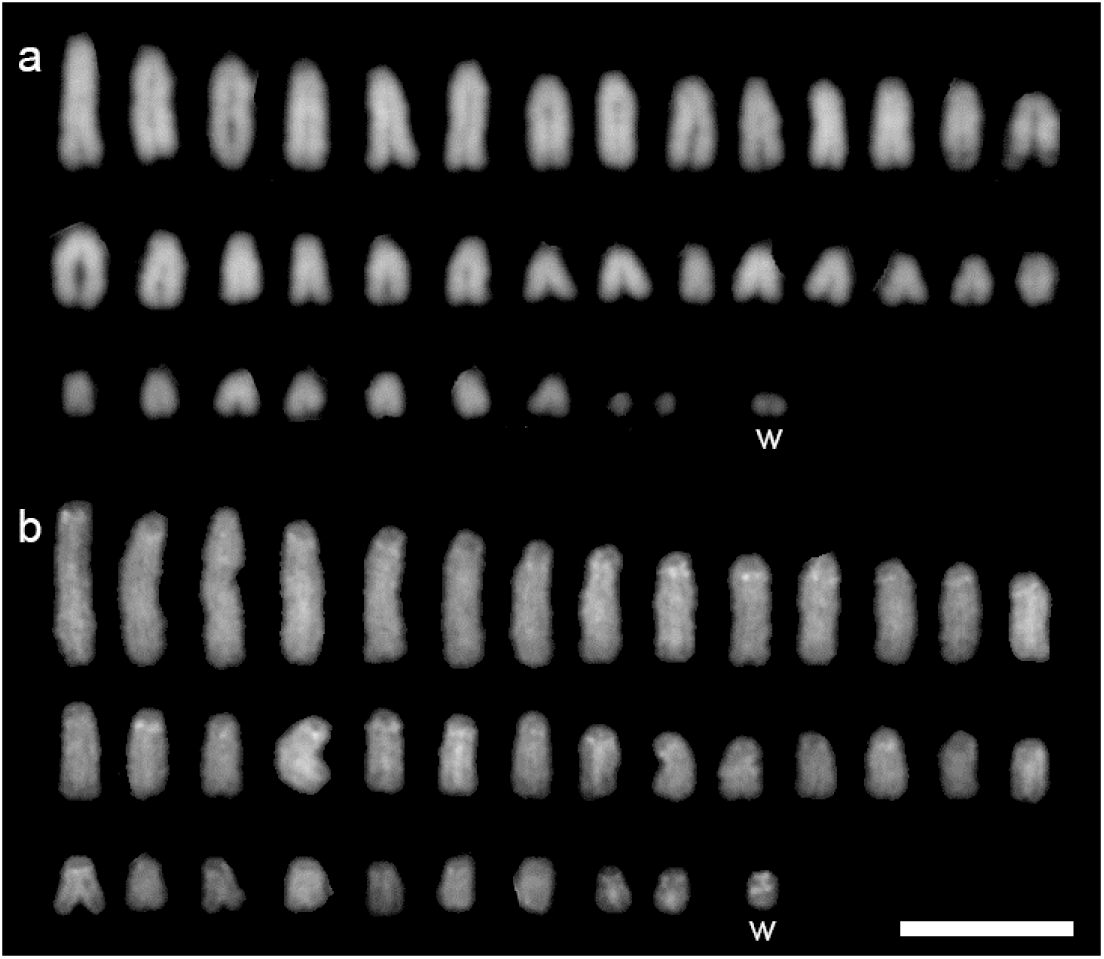
Karyotypes of *L. strigata* (a) and *L. agilis* (b). DAPI staining. Scale bar represents 10 μm.

**Fig. 2.**
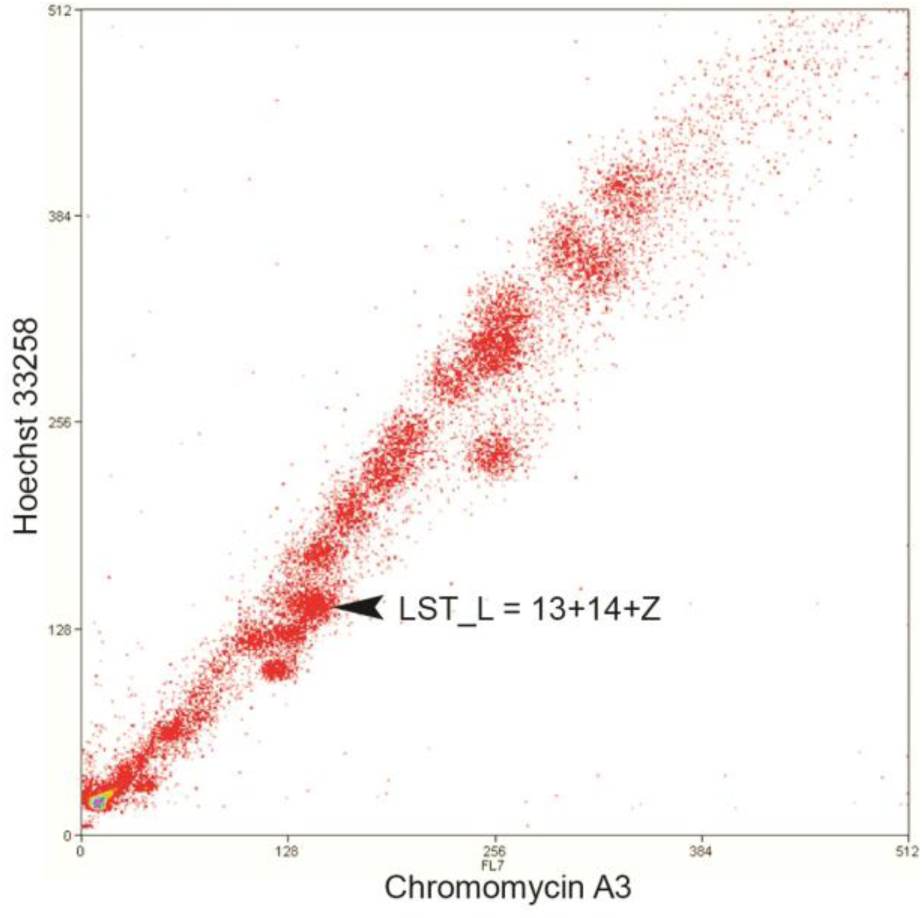
Flow-sorted karyotype of *L. strigata.*

**Fig. 3.**
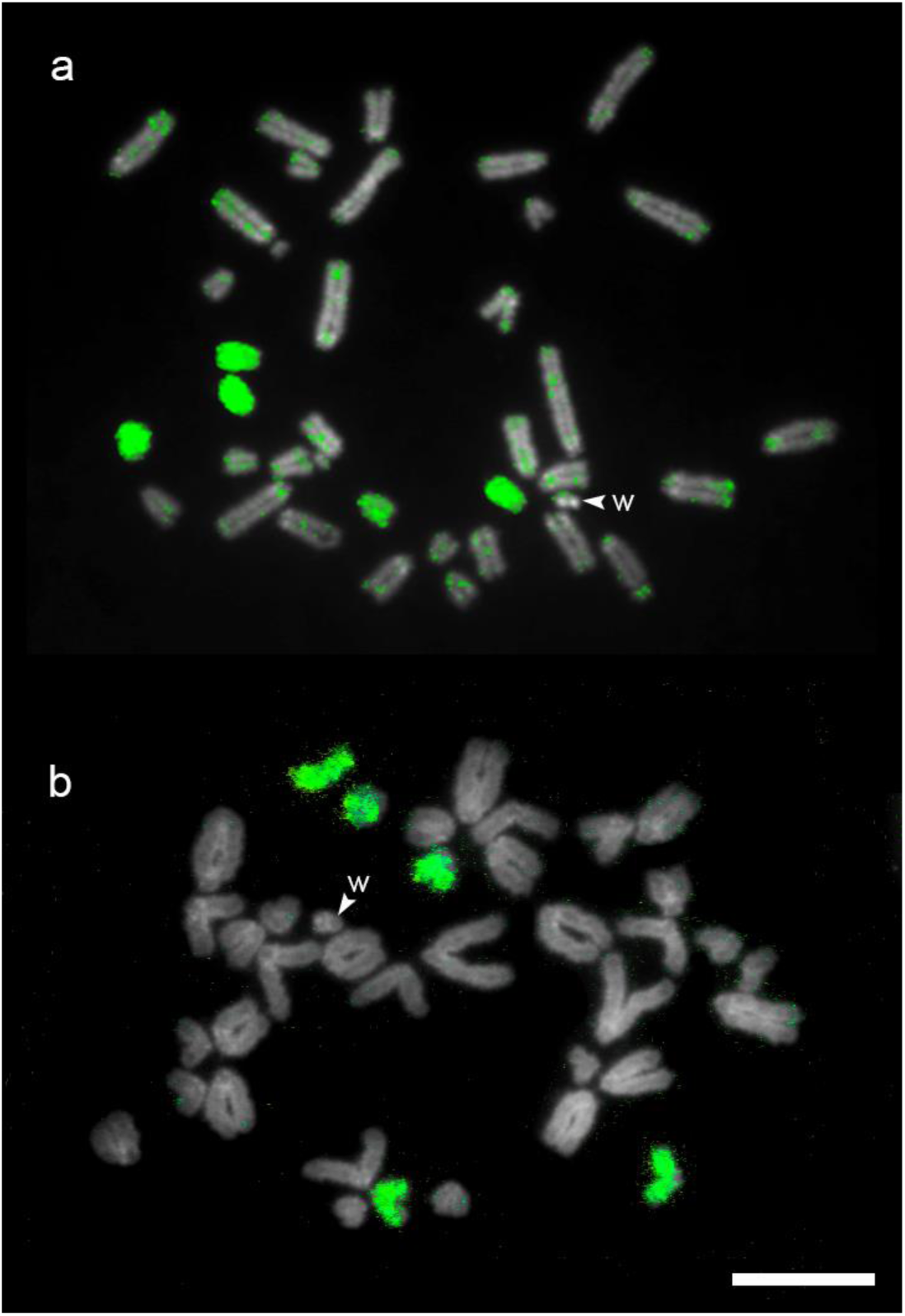
FISH with the LST_L probe (green) on *L. strigata* (a) and *L. agilis* (b). Scale bar represents 10 μm.

The karyotype of *L. agilis* also consisted of 38 chromosomes, including the heteromorphic W chromosome, the smallest element of the karyotype (Fig. 1b). This result agrees with the previous analyses of the karyotype of *L. agilis* [De Smet, 1981]. FISH with the LST_L probe showed hybridization with five small macrochromosomes, as for the same-species hybridization (Fig. 3b). FISH with the telomeric probe showed the accumulation of the telomeric repeat on the W chromosome, as shown previously for this species [Matsubara et al., 2015; Giovannotti et al., 2018] (Fig. 4).

**Fig. 4.**
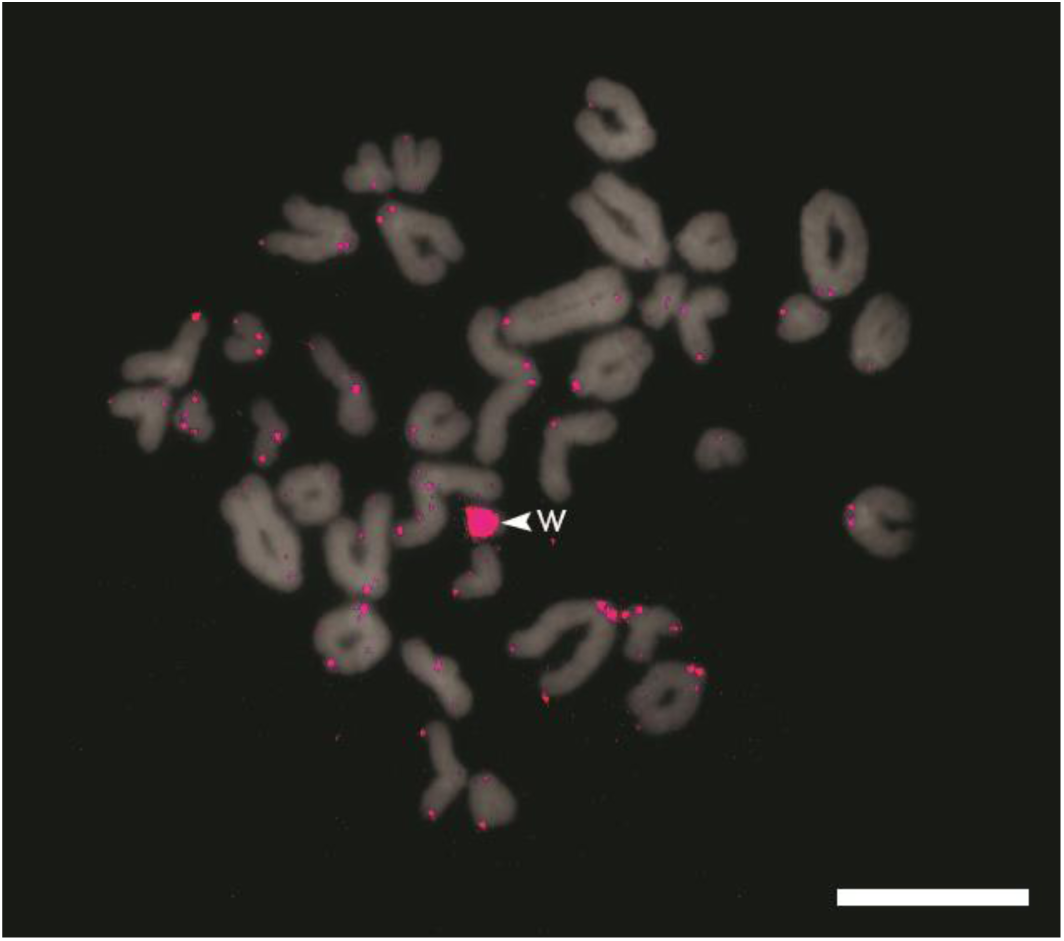
FISH with the telomeric probe (red) on *L. agilis*. Scale bar represents 10 μm.

## Discussion

Our results do not confirm the identification of LAG5 as a sex chromosome. This resolves the five-year long debate concerning the sex chromosomes of the sand lizard.

The fact that the LST_L probe painted five chromosomes in *L. agilis* as well as in *L. strigata* indicates that these two species share the same sex chromosome system. If *L. agilis* had experienced a hypothetical sex chromosome turnover, leading to the formation of neo-sex chromosomes derived from the LAG5 pair, and the original lacertid Z chromosome had returned to the autosomal state, as shown for sex chromosome turnovers in the genus *Paroedura* (Gekkonidae) [Koubová et al., 2014], the LST_L probe would have hybridized with six chromosomes (three homologous pairs). The hybridization of the LST_L probe with five chromosomes was observed in multiple complete metaphases, which confirms the credibility of our results.

The possibility that these two species share the same sex chromosome turnover is excluded by the size of the chromosomes which constitute the LST_L probe: they belong to the fraction of small chromosomes, whereas LAG5/LST5 are nearly twice as large. The small size of the Z chromosome in *L. agilis* and *L. strigata* fits well with the results of Rovatsos et al. [2016, 2019] and Andrade et al. [2019], which showed that the lacertid Z originates from two fused microchromosomes of ancestral squamates.

There could be still a possibility that the sex chromosome turnover is not universal for *L. agilis*, but it is specific to the lineage studied by Srikulnath et al. [2014] and Matsubara et al. [2015]: the nominative subspecies *L. agilis agilis*. However, the features of the W chromosome of the lizards which were studied in these works provide evidence against this explanation.

First, in case of such a recent turnover, the Z and W chromosomes would be expected to be homomorphic, whereas the W chromosome in *L. agilis agilis* is highly divergent from the putative Z chromosome [Srikulnath et al., 2014; Matsubara et al., 2015].

Second, the relative size and telomeric repeat accumulation of the *L. agilis agilis* W chromosome correspond well with the features of the W chromosome of *L. agilis exigua* that were observed in this study. Furthermore, W chromosomes of similar size and repeat content are characteristic for other species of *Lacerta*: all studied *Lacerta* species including *L. agilis* have telomeric repeat accumulation on the W chromosome, and share a monophyletic W-specific subfamily of the IMO-TaqI satellite DNA [Giovannotti et al., 2018]. Considering the fact that the sequence content of the W chromosomes in the related genera *Iberolacerta* and *Timon* is different from that of *Lacerta*, as shown by CGH [Rojo Oróns, 2015] and that W-specific repetitive DNA can also be different in different populations of the same species [Giovannotti et al., 2017], the sharing of the same repetitive DNAs by the W chromosomes of different *Lacerta* species seems to indicate stability and homology for the ZZ/ZW system within this genus.

Thus, we conclude that the sex chromosomes of *L. agilis* specimens studied by Srikulnath et al. [2014] are most probably not different from the sex chromosomes of *L. agilis* individuals studied in the current work. Therefore, it is unlikely that LAG5 is the Z chromosome in this species. Recently, Rovatsos et al. [2019] doubled the number of lacertid species included in their analysis, and found no confirmation for several other putative cases of sex chromosome turnovers in some lacertids. Our results agree with this conclusion and reinforce the idea of stability of the lacertid ZZ/ZW sex chromosome system.

The library of flow-sorted chromosome probes of *L. strigata* is a powerful tool to study chromosome evolution in Lacertidae. It may be further used to test the sex chromosome homology and identify chromosomal rearrangements in other species of lacertids, which were previously not studied using the molecular cytogenetic methods.

## Acknowledgement

We thank S. O. Baturin and A. I. Stekleneva (Institute of Cytology and Genetics) for help in obtaining the specimen of *L. agilis exigua*. We thank the Microscopic Center of the Siberian Branch of the Russian Academy of Sciences for granting access to microscopic equipment.

## Statement of ethics

All manipulations with live animals and euthanasia were approved by the Institute of Molecular and Cellular Biology Ethics Committee (statement #01/18 from 05.03.2018).

## Author contributions

AL, VT, and PB designed the study. AL produced the figures and wrote the initial draft of the manuscript. MG and SR prepared the cell cultures. JP and MFS performed flow sorting. JP and DA performed FISH. All authors participated in writing and editing the manuscript.

## References

Alföldi J, Di Palma F, Grabherr M, Williams C, Kong L, et al.: The genome of the green anole lizard and a comparative analysis with birds and mammals. Nature 477: 587–591 (2011).

Andrade P, Pinho C, i de Lanuza GP, Afonso S, Brejcha J, et al: Regulatory changes in pterin and carotenoid genes underlie balanced color polymorphisms in the wall lizard. Proceedings of the National Academy of Sciences 116(12): 5633–5642 (2019).

Capriglione T, Olmo E, Odierna G, Kupriyanova LA: Mechanisms of differentiation in the sex chromosomes of some Lacertidae. Amphibia-Reptilia 15: 1–8 (1994).

De Smet WHO: Description of the orcein stained karyotypes of 36 lizard species (Lacertilia, Reptilia) belonging to the families Teiidae, Scincidae, Lacertidae, Cordylidae and Varanidae (Autarchoglossa). Acta Zool Pathol Antverpiensia 76:73–118 (1981).

Deakin JE, Ezaz T: Understanding the Evolution of Reptile Chromosomes through Applications of Combined Cytogenetics and Genomics Approaches. Cytogenetic and Genome Research 157(1-2): 7–20 (2019).

Gamble T: A review of sex determining mechanisms in geckos (Gekkota: Squamata). Sexual Development 4(1-2): 88–103 (2010).

Gamble T, Geneva AJ, Glor RE, Zarkower D: *Anolis* sex chromosomes are derived from a single ancestral pair. Evolution 68: 1027–1041 (2014).

Giovannotti M, Nisi Cerioni P, Slimani T.,Splendiani A, Paoletti A, et al: Cytogenetic characterization of a population of *Acanthodactylus lineomaculatus* Duméril and Bibron, 1839 (Reptilia, Lacertidae) from South-western Morocco and insights into sex chromosome evolution. Cytogenetic and Genome Research 153: 86–95 (2017).

Giovannotti M, Nisi Cerioni P, Rojo V, Olmo E, Slimani T, et al: Characterization of a satellite DNA in the genera *Lacerta* and *Timon* (Reptilia, Lacertidae) and its role in the differentiation of the W chromosome. Journal of Experimental Zoology Part B Molecular and Developmental Evolution 330: 83–95 (2018).

Giovannotti M, Trifonov VA, Paoletti A, Kichigin IG, O’Brien PCM, et al.: New insights into sex chromosome evolution in anole lizards (Reptilia, Dactyloidae). Chromosoma 126(2): 245–260 (2017).

Graphodatsky AS, Sablina OV, Meyer MN, Malikov VG, Isakova EA, et al.: Comparative cytogenetics of hamsters of the genus *Calomyscus*. Cytogenetics and Cell Genetics 88: 296–304 (2000).

Graphodatsky AS, Yang F, O’Brien PC, Perelman P, Milne BS, et al.: Phylogenetic implications of the 38 putative ancestral chromosome segments for four canid species. Cytogenetics and Cell Genetics 92: 243–247 (2001).

Holleley CE, O’Meally D, Sarre, SD, Graves JAM, Ezaz T, et al.: Sex reversal triggers the rapid transition from genetic to temperature-dependent sex. Nature 523(7558): 79–82 (2015).

Ijdo JW, Wells RA, Baldini A, Reeders ST: Improved telomere detection using a telomere repeat probe (TTAGGG)n generated by PCR. Nucleic Acids Research 19(17): 4780 (1991).

Ivanov VG, Fedorova TA: Sex heteromorphism of chromosomes in *Lacerta strigata* Eichwald. Tsitologiya 12: 1582–1585 (1970).

Kichigin IG, Giovannotti M, Makunin AI, Ng BL, Kabilov ML, et al.: Evolutionary dynamics of *Anolis* sex chromosomes revealed by sequencing of flow sorting-derived microchromosome-specific DNA. Molecular Genetics and Genomics 291(5): 1955–1966 (2016).

Koubová M, Pokorná MJ, Rovatsos M, Farkačová K, Altmanová M, Kratochvíl L: Sex determination in Madagascar geckos of the genus *Paroedura* (Squamata: Gekkonidae): are differentiated sex chromosomes indeed so evolutionary stable? Chromosome Research 22: 441– 452 (2014).

Liehr T, Kreskowski K, Ziegler M, Piaszinski K, Rittscher K: The Standard FISH Procedure. In: Liehr T (Ed.) Fluorescence in Situ Hybridization (FISH). Berlin, 109–118 (2017).

Lisachov AP, Makunin AI, Giovannotti M, Pereira JC, Druzhkova AS, et al. Genetic content of the neo-sex chromosomes in *Ctenonotus* and *Norops* (Squamata, Dactyloidae) and degeneration of the Y chromosome as revealed by high-throughput sequencing of individual chromosomes. Cytogenetic and Genome Research 157(1-2):115–122 (2019).

Matsubara K, Sarre SD, Georges A, Matsuda Y, Graves JAM, Ezaz T: Highly differentiated ZW sex microchromosomes in the Australian *Varanus* species evolved through rapid amplification of repetitive sequences. PLoS One 9(4): e95226 (2014).

Matsubara K, Tarui H, Toriba M, Yamada K, Nishida-Umehara C, et al.: Evidence for different origin of sex chromosomes in snakes, birds, and mammals and step-wise differentiation of snake sex chromosomes. Proceedings of the National Academy of Sciences 103(48): 18190–18195 (2006).

Matsubara K, Uno Y, Srikulnath K, Matsuda Y, Miller E, Olsson M: No interstitial telomeres on autosomes but remarkable amplification of telomeric repeats on the W sex chromosome in the sand lizard (*Lacerta agilis*). Journal of Heredity 106: 753–757 (2015).

Matveevsky S, Kolomiets O, Bogdanov A, Hakhverdyan M, Bakloushinskaya I: Chromosomal evolution in mole voles *Ellobius* (Cricetidae, Rodentia): Bizarre sex chromosomes, variable autosomes and meiosis. Genes 8(11): 306 (2017).

Nielsen SV, Guzmán-Méndez IA, Gamble T, Blumer M, Pinto BJ, et al.: (2019). Escaping the evolutionary trap? Sex chromosome turnover in basilisks and related lizards (Corytophanidae: Squamata). Biol Lett 15(10): 20190498 (2019).

Ogata M, Lambert M, Ezaz T, Miura I: Reconstruction of female heterogamety from admixture of XX-XY and ZZ-ZW sex-chromosome systems within a frog species. Molecular Ecology 27(20): 4078–4089 (2018).

Pokorná M, Kratochvíl L: Phylogeny of sex-determining mechanisms in squamate reptiles: are sex chromosomes an evolutionary trap? Zoological Journal of the Linnean Society, 156(1): 168–183 (2009).

Pokorná MJ, Kratochvíl L: What was the ancestral sex-determining mechanism in amniote vertebrates? Biological Reviews 91(1): 1–12 (2016).

Rojo Oróns V: Cytogenetic and molecular characterization of lacertid lizard species from the Iberian Peninsula. Doctoral thesis, Universidade da Coruña (2015).

Romanenko SA, Biltueva LS, Serdyukova NA, Kulemzina AI, Beklemisheva VR, et al: Segmental paleotetraploidy revealed in sterlet (*Acipenser ruthenus*) genome by chromosome painting. Molecular Cytogenetics 8: 90 (2015).

Rovatsos M, Pokorná M, Altmanová M, Kratochvíl L: Cretaceous park of sex determination: sex chromosomes are conserved across iguanas. Biol Lett 10(3): 20131093 (2014).

Rovatsos M, Rehák I, Velenský P, Kratochvíl L: Shared ancient sex chromosomes in varanids, beaded lizards, and alligator lizards. Molecular biology and evolution 36(6): 1113–1120 (2019a).

Rovatsos M, Vukić J, Altmanová M, Johnson Pokorná M, Moravec J, Kratochvíl L: Conservation of sex chromosomes in lacertid lizards. Molecular Ecology 25(13): 3120–3126 (2016).

Rovatsos M, Vukić J, Lymberakis P, Kratochvíl L: Evolutionary stability of sex chromosomes in snakes. Proc. R. Soc. B 282(1821): 20151992 (2015).

Rovatsos M, Vukić J, Mrugała A, Suwala G, Lymberakis P, Kratochvíl L: Little evidence for switches to environmental sex determination and turnover of sex chromosomes in lacertid lizards. Scientific Reports 9(1): 7832 (2019b).

Srikulnath K, Matsubara K, Uno Y, Nishida C, Olsson M, Matsuda Y: Identification of the linkage group of the Z sex chromosomes of the sand lizard (*Lacerta agilis*, Lacertidae) and elucidation of karyotype evolution in lacertid lizards. Chromosoma 123: 563–575 (2014).

Stanyon R, Galleni L: A rapid fibroblast culture technique for high resolution karyotypes. Bolletino di Zoologia 58: 81–83 (1991).

Telenius H, Ponder BA, Tunnacliffe A, Pelmear AH, Carter NP, et al.: Cytogenetic analysis by chromosome painting using dop-pcr amplified flow-sorted chromosomes. Genes, Chromosomes and Cancer 4(3): 257–263 (1992).

Uno Y, Nishida C, Tarui H, Ishishita S, Takagi C, et al.: Inference of the protokaryotypes of amniotes and tetrapods and the evolutionary processes of microchromosomes from comparative gene mapping. PLoS One 7(12): e53027 (2012).

Vicoso B, Emerson JJ, Zektser Y, Mahajan S, Bachtrog D: Comparative sex chromosome genomics in snakes: differentiation, evolutionary strata, and lack of global dosage compensation. PLoS Biol 11(8): e1001643 (2013).

Viets BE, Tousignant A, Ewert MA, Nelson CE, Crews D: Temperature-dependent sex determination in the leopard gecko, *Eublepharis macularius*. Journal of Experimental Zoology 265(6): 679–683 (1993).

Yang F, Carter NP, Shi L, Ferguson-Smith MA: A comparative study of karyotypes of muntjacs by chromosome painting. Chromosoma 103: 642–652 (1995).

Yang F, O’Brien PC, Milne BS, Graphodatsky AS, Solanky N, et al: A complete comparative chromosome map for the dog, red fox, and human and its integration with canine genetic map. Genomics 62: 189–202 (1999).

